# Soil disturbance affects plant growth via soil microbial community shifts

**DOI:** 10.1101/2020.10.16.343053

**Authors:** Taylor J. Seitz, Ursel M. E. Schütte, Devin M. Drown

## Abstract

Recent advances in climate research have discovered that permafrost is particularly vulnerable to the changes occurring in the atmosphere and climate, especially in Alaska where 85% of the land is underlain by mostly discontinuous permafrost. As permafrost thaws, research has shown that natural and anthropogenic soil disturbance causes microbial communities to undergo shifts in membership composition and biomass, as well as in functional diversity. Boreal forests are home to many plants that are integral to the subsistence diets of many Alaska Native communities. Yet, it is unclear how the observed shifts in soil microbes can affect above ground plant communities that are relied on as a major source of food. In this study, we tested the hypothesis that microbial communities associated with permafrost thaw affect plant growth by growing five plant species found in Boreal forests and Tundra ecosystems, including low-bush cranberry and bog blueberry, with microbial communities from the active layer soils of a permafrost thaw gradient. We found that plant growth was significantly affected by the microbial soil inoculants. Plants inoculated with communities from above thawing permafrost showed decreased growth compared to plants inoculated with microbes from undisturbed soils. We used metagenomic sequencing to determine that microbial communities from disturbed soils above thawing permafrost have differences in taxonomy from microbial communities in undisturbed soils above intact permafrost. The combination of these results indicates that a decrease in plant growth can be linked to soil disturbance driven changes in microbial community membership and abundance. These data contribute to an understanding of how microbial communities can be affected by soil disturbance and climate change, and how those community shifts can further influence plant growth in Boreal forests and more broadly, ecosystem health.

## 1 Introduction

With nearly 85% of the land in Alaska underlain with discontinuous permafrost, Alaska is particularly vulnerable to large-scale ecosystem changes due to climate change-driven permafrost thaw and resulting soil disturbance (Department of Natural Resources, 2020). As foundational abiotic factors such as nutrient availability and soil hydrology begin to change, they in turn affect ecosystem processes such as succession and productivity, that regulate plant and microbial communities present in the disturbed soils (Schuur et al., 2018). Recent research has turned to exploring the effects of climate change on microbial communities residing in soil and permafrost (Jansson and Hofmockel, 2018). As permafrost thaws and soil disturbance events occur, microbial communities undergo shifts in membership composition and biomass, as well as in functional diversity (Schuur et al., 2015; Mackelprang et al., 2011; Schütte et al., 2019). As these smaller scale changes develop, it is important that we build a better understanding of how microbial communities in northern latitude soils affect larger scale processes such as plant growth.

Permafrost is frozen ground (rock, ice, soil) that remains at less than 0° C for more than two years. Permafrost underlies approximately 26% of terrestrial ecosystems globally and is a critical structural element that holds large potential for altering ecosystem composition through thawing and melting of ice wedges (Schuur et al., 2018). It has been estimated that the carbon stored in permafrost accounts for more than 2.5 times as much as the atmospheric carbon pool (Schuur et al., 2007; Schuur et al., 2015). As permafrost thaws, previously frozen organic matter, nutrients, water, and microbes are exposed and reintroduced into actively cycling pools of carbon and nutrients (Chapin et al., 2006). Plant roots interact within the active layer of soil which sits above the permafrost and undergoes seasonal cycles of freezing and thawing (Schuur et al., 2015; Christensen et al., 2004). The depth of the active layer defines what nutrients, rooting space, and water are available to plants which in turn can affect processes such as community succession and species composition.

The atmospheric release of carbon trapped in soils following permafrost thaw is mediated by soil bacterial and fungal respiration. These organisms have been surviving in the active layer and frozen soil below for millennia, either frozen or respiring at extremely low rates, using the frozen organic matter as an energy source (Schütte et al., 2019; Tripathi et al., 2018). As the permafrost thaws, soil microbes exit dormancy and microbial communities gain access to large pools of previously frozen carbon and nutrients (Burkert et al., 2019). As the activity of the microbial communities increases with a warming climate, microbial effects may be more complicated than merely releasing greenhouse gases into the environment through respiration. These microbes can also cause alterations and feedback to the ecosystem nutrient cycling and net primary production (du Toit, 2018; Natali et al., 2012).

Many studies focused in the arctic and sub-arctic have demonstrated that soil microbes are important drivers of plant succession, relative abundance, and productivity (Mangan et al., 2010; Natali et al., 2012; Schuur et al., 2007). As physical and chemical properties of soil such as temperature and pH are changing, the composition of microbial communities present in the active layer changes as well. Permafrost thaw could induce now active microbes to transform newly available organic matter into gaseous products such as CO_2_ and CH_4_, and metabolites that are now accessible to plants and other organisms living in active layer soils (Graham et al., 2012). With the availability of newly released nutrients and the production of metabolites, soil microbes then have the potential to shape above ground plant communities by mediating and partitioning soil resources (Ho et al., 2017).

Previous research has found evidence that the composition, diversity, and biomass of microbial communities can become altered in locations that are undergoing permafrost thaw and disturbance events such as thermokarst formation (Mackelprang et al., 2011; Monteux et al., 2019; Schurr et al., 2007). These shifts in microbial community characteristics suggest that microbes affected by thawing permafrost may in turn drive alterations to the plant community in associated soils. Here we investigate how active layer soil microbial communities affect above ground plant communities, indicating that microbes play a role in climate change driven alterations to Alaskan plant communities. We hypothesized that microbial communities from disturbed soils associated with greater permafrost thaw would lower levels of plant growth compared to plants inoculated with microbes from less disturbed soils. We tested this hypothesis by conducting a plant growth experiment on boreal plant species inoculated with microbes from soils with different degrees of permafrost thaw and assessed soil microbial community composition.

## 2 Materials and Methods

### 2.1 Sample Site Description and Soil Collection

The Fairbanks Permafrost Experiment Station (FPES) is located in interior Alaska (64.877°N, 147.670°W) and is part of the US Army Corps of Engineers Cold Regions Research and Engineering Laboratory (CRREL). FPES is divided into three Linell plots (Linell, 1973; Douglas et al., 2008), each 3,721 m^2^: undisturbed (UD), semi disturbed (SD), and most disturbed (MD) (SFigure 1). When these three plots were created in 1946, the first plot, UD, was left untouched. The second plot (SD) was cleared of trees and above ground vegetation by hand, but the roots and other organic material were left intact. The third plot (MD) was stripped of all vegetation and surface level organic material down to mineral soil. Over the next 25 years, total stripping (MD) led to permafrost thaw and degradation to 6.7 m below surface level, and partial stripping (SD) led to thaw 4.7 m below surface level. These plots were created to simulate soil disturbance events and to then identify potential ecological effects.

Each plot is representative of the subarctic taiga forest. The undisturbed plot is covered by a dense black spruce stand (*Picea mariana*) with intermittent white spruce (*Picea glauca*). Its understory is dominated by Labrador tea (*Rhododendron groenlandicum*) and low-bush cranberry (*Vaccinium vitis-idaea*) with a ground cover of continuous feather and *Sphagnum* moss and lichen. The surface soils are moderately moist and can be classified as mesic, with an average soil organic layer thickness of 35 cm and a consistent thaw depth of 85 cm (Johnstone et al., 2008; Douglas et al., 2008). The semi-disturbed plot is covered by a mixed stand of Alaskan birch (*Betula neoalaskana*), willow (*Salix sp.*), black spruce, and white spruce, with a developing understory of moss and shrubs. The trees at the semi-disturbed plot are taller than those in the most disturbed. The most disturbed plot is covered by a mixed stand of Alaskan birch, willow, and young black spruce. The understory is a mixture of moss and grass (Douglas et al., 2008).

We collected 48 soil cores from FPES on 28 May 2018, 16 individual cores at each FPES treatment level. Using an established grid layout of FPES, we took cores from four selected quadrats per plot to demonstrate the fine-scale heterogeneity of the sample site. At each quadrat we took four samples at the corners, approximately 1 m apart. We utilized a sterile technique to sample soil and at each sampling point, the top layer of moss and vegetation was removed. We used a soil corer (4.5 cm diameter by 10 cm height) to collect the top 10 cm of soil, which were stored in a cooler throughout the duration of sample collection. We then transported the cores back to the lab where each was homogenized and stored at 4° C to be used 10 days later for soil inoculants and between 7 to 45 days later for DNA extractions.

### 2.2 Greenhouse Experimental Design and Sampling

To test how the sampled microbial communities affect plant growth, we grew all plants in a climate controlled greenhouse, within autoclaved, nutrient-poor soils containing soil microbial inoculum. The soil microbial inoculants were obtained from FPES for the three treatment levels: UD, SD, and MD. We also included a sterile inoculant (ST), an autoclaved mixture of UD, SD, and MD soils.

We conducted the growth experiment in the Institute of Arctic Biology Greenhouse at the University of Alaska Fairbanks (UAF), consisting of five plant species: *Vaccinium vitis-idaea* (low-bush cranberry), *Vaccinium uliginosum* (bog blueberry); *Picea mariana* (black spruce); *Ledum groenlandicum* (Labrador tea); and *Chamerion angustifolium* (fireweed), hereafter all plants will be referred to by their colloquial names. We obtained all seeds from Alaskan sources, and we surface-sterilized them prior to planting. We placed five to ten seeds of one plant type into a pot (SC10 Cone-tainers, Ray Leach; USA) filled with a mixture of densely packed sterile vermiculite and Canadian peat (ProMix), with 1.5 g of soil containing the treatment microbial soil inoculant (Figure 1). Sixty-four pots were planted for each plant type, with 16 unique soil inoculants per each of four treatments, for a total of 320 pots. We randomized all pots across seven RL98 trays (Ray Leach, USA) to reduce possible bias from temperature or light gradients within the greenhouse. Plants were maintained under 12hr light cycles and watered once daily.

**Figure 1.**
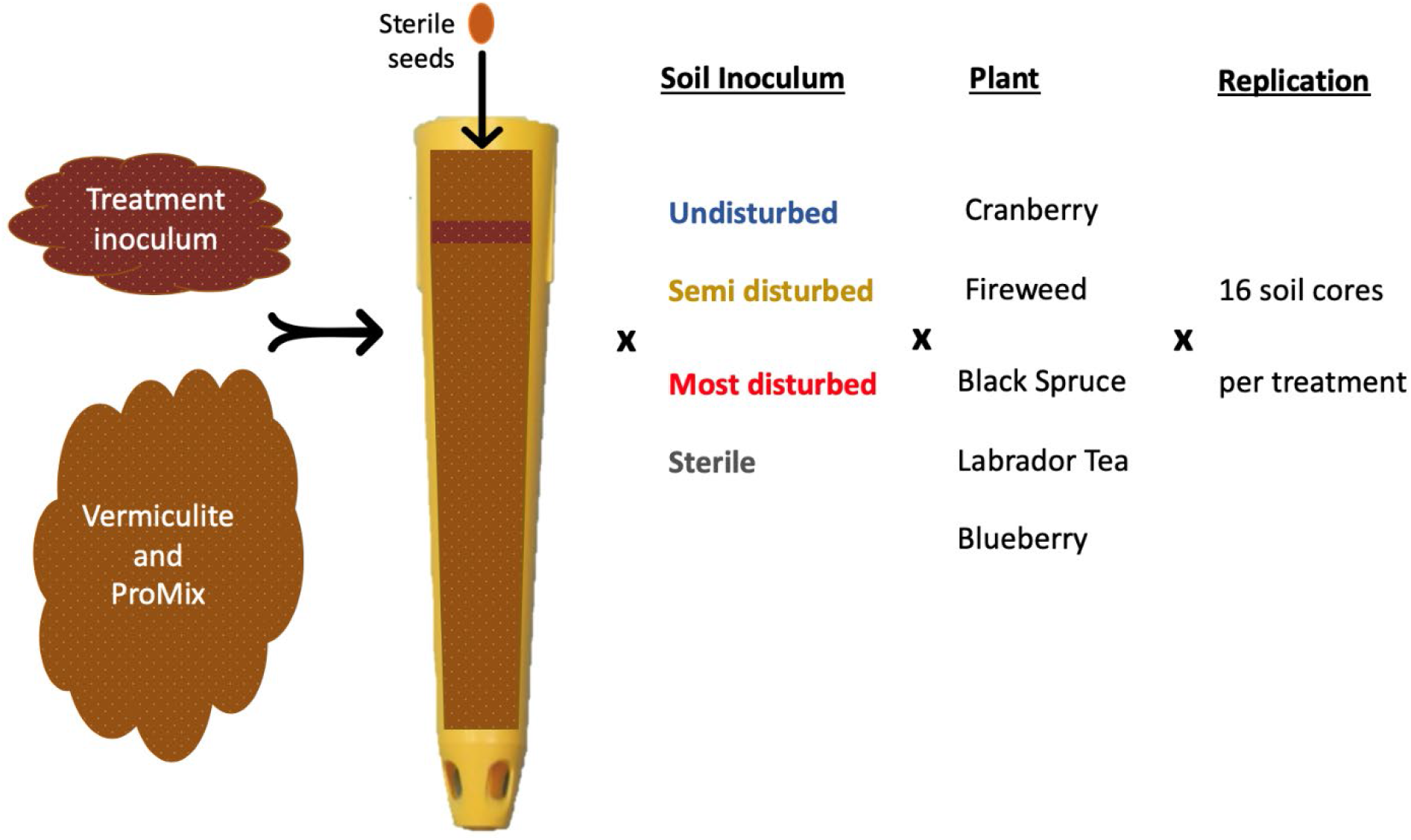
Overview of experimental design for the greenhouse growth experiment.

Following the first month of growth past germination, we measured the plants biweekly for the first three months, recording their leaf count and height to the nearest millimeter. We stopped black spruce needle counts in November 2018, when needle counts exceeded 200 and we were unable to accurately measure. By October 2018, all the fireweed (n = 61) had started to decline in height measurements, so on 10 October 2018, we clipped above ground living plant material at the soil surface and then oven dried plants 55 °C to a constant mass before determining biomass produced. The bog blueberry, black spruce, Labrador tea, and low-bush cranberry continued to grow and be measured monthly. We then harvested above ground biomass per plant type in March 2020, and below ground biomass in April 2020. We oven dried plants at 55° C to a constant mass before weighing.

### 2.3 Statistical Analysis

All statistical analyses were performed in R version 3.5.2 (R Core Team, 2014). In order to test the effects of the microbial inoculant treatments during the plant growth experiment, we used the Shapiro-Wilk test and Levene’s test to check ANOVA assumptions. We log-transformed growth measurements (leaf count, height, biomass) per plant type if it improved normality and reduced heteroscedasticity. We used a one-way ANOVA (Car R package, version 3.0.3) to evaluate the significance of soil inoculant on each plant growth measurement within each plant type. We used a Tukey HSD (Agricolae R package, Mendiburu, 2009, version 1.3.2) post-hoc test to identify significant effects of specific inoculant treatments on growth within each plant type. We visualized differences using the R package ggplot2 (version 3.2.1).

### 2.4 DNA Extractions and Metagenomic Sequencing

In order to analyze the microbial communities used to inoculate plants in the growth experiment, we performed shotgun metagenomic sequencing on each individual core. To do this, we extracted and purified total genomic DNA from approximately 250 mg of soil per homogenized soil core using the DNeasy PowerSoil kit (Qiagen; Germany) following manufacturer instructions. We quantified the yield and the quality of the DNA extracted using a NanoDrop One spectrophotometer (Thermo Scientific; USA) and a Qubit (Thermo Scientific; USA). Following DNA extractions, we randomly divided the 48 cores into four sequencing runs. We prepared the DNA sequencing library using the Oxford Nanopore Technologies Ligation Kit with Native barcodes to multiplex 12 samples (SQK-LSK109, ONT; UK). We diluted sample DNA to 400 ng for input and followed the kit according to the manufacturer instructions. We then sequenced DNA using a MinION and R9.4.1 and R9.5 flow-cells (FLO-MIN106) (Table S1). The sequencing runs each lasted 48 hours.

### 2.5 Metagenomic Analyses

We base called the raw data using Guppy v3.6.1 (ONT) specifying the high accuracy model (Table 1). We then demultiplexed samples with the Guppy barcoder using parameters to discard sequences with middle adapters and to trim barcodes. We used Filtlong Filtlong v0.2.0 (https://github.com/rrwick/Filtlong) to control for length (≥ 50 bp) and quality (Q) score (≥ 10). Following quality control, we detected taxa with a *k*-mer based approach using Kraken 2 (v.2.0.9-beta; Wood et al., 2019) and subsequently estimated abundance using Bracken (v.2.6.0; Lu et al., 2017). We used a reference database built using complete genomes from RefSeq for the bacterial, archeal, and viral domains.

We merged sample Bracken reports and then generated heat maps with pheatmap (version 1.0.12; Kolde, 2019) using relative abundances scaled by z-score. In order to identify biomarkers within the microbial communities we used the linear discriminant analysis (LDA) effect size (LEfSe) method (Segata et al., 2011). To accomplish this, we uploaded our Bracken results to the Galaxy web platform (https://galaxyproject.org/; Afgan et al., 2018) where we then used the Huttenhower Lab workflow (https://huttenhower.sph.harvard.edu/galaxy/) for LEfSe analysis. We used an LDA threshold of 2.0 and significance α of 0.05 to detect biomarkers.

## 3 Results

### 3.1 Plant Growth Experiment

Our results indicate that soil microbial communities from across the thaw gradient differentially affected plant productivity (Figures 2–6; Tables S3-S8). Most plant types responded negatively when grown in soils inoculated with microbial communities from the most disturbed FPES soil compared to either inoculants from semi disturbed or undisturbed soils, or the sterile treatment (Figure S2; Table S8).

**Figure 2.**
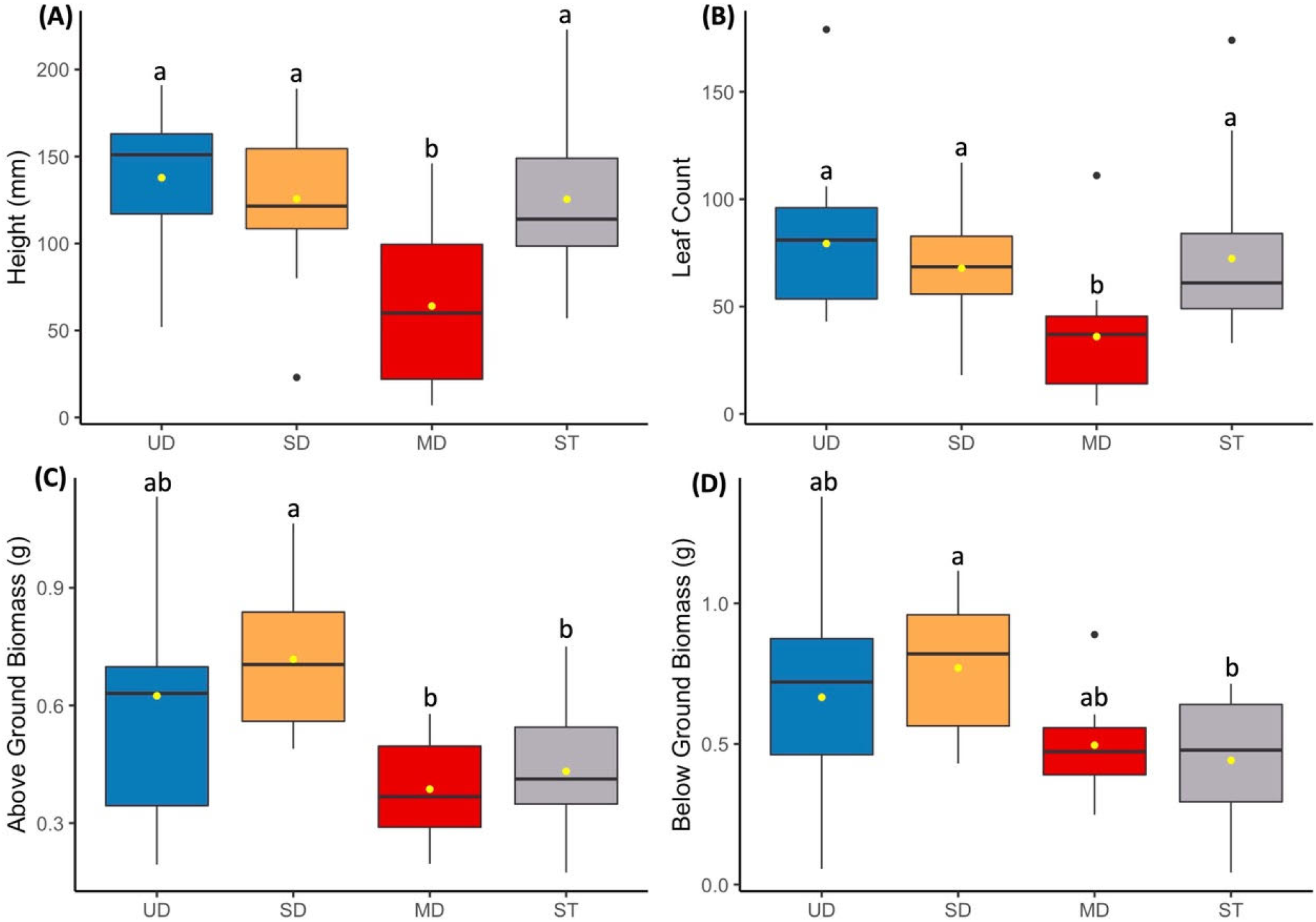
Bog blueberry plant height (mm) (A) and leaf count (B) at 184 days since planting. Above ground (C) and below ground (D) biomass after 1.5 years since planting. Boxplots represent median, and the upper and lower quartiles. Yellow circles represent mean value for each group. Tukey HSD post-hoc comparisons are denoted with lowercase letters above boxplots.

#### Bog Blueberry

Bog blueberry had reduced growth in height and leaf count when grown in soils inoculated with microbial communities from the MD treatment site compared to plants growth with inoculants from UD, SD, or ST soil (Figure 2). This trend was consistent across height and leaf count starting at the third through the final measurement point (Figure S2). While the mean height and leaf count of bog blueberry grown in MD treatment was significantly smaller compared to bog blueberry grown under the SD, UD, or ST treatments (Table S3), there was no significant difference between height or leaf number for bog blueberry plants grown in UD, SD, or ST treatments. While the source of inoculant was a significant factor in above ground and below ground biomass, the responses did not follow the same trends as height and leaf count. For above ground biomass the post hoc Tukey HSD test showed that SD plants had more mass on average and differed significantly from MD and ST, SD plants did not differ significantly in biomass production (Table S8).

#### Low-bush Cranberry

Low-bush cranberry showed the same trends to bog blueberry when grown in soils from the most disturbed treatment site (Figure 3). Low-bush cranberry height and leaf count significantly decreased when grown with inoculant from the MD site compared to when grown with inoculant from the SD or UD sites, or with ST inoculant (Table S4, Table S8). There was no significant difference in either height or leaf count between low-bush cranberry grown in UD, SD, or ST treatments. Above ground biomass increased when plants were grown with soil from the SD compared to plants inoculated with UD, MD, and ST treatments. Low-bush cranberry below ground biomass did not differ significantly depending on the soil inoculant treatment.

**Figure 3.**
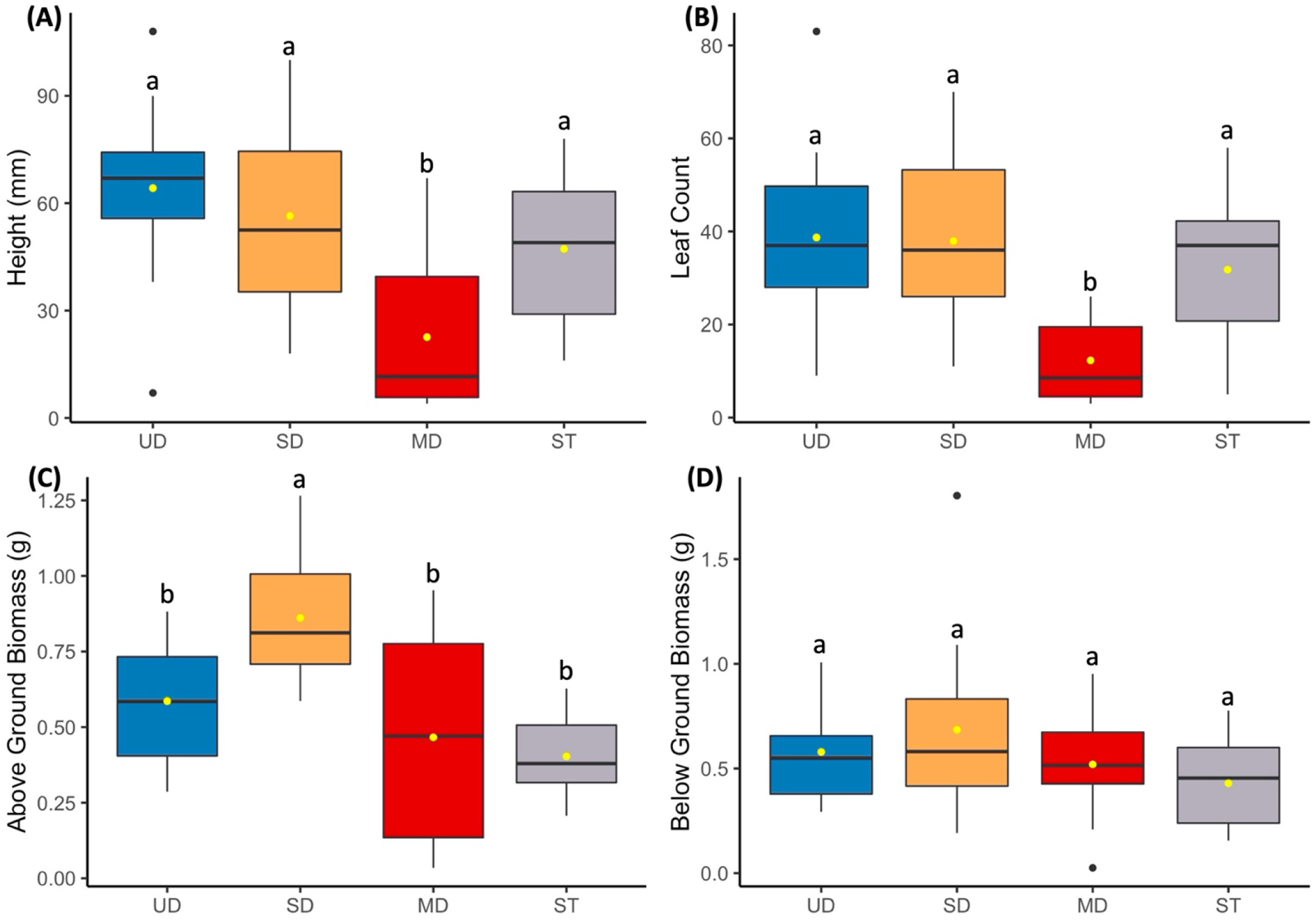
Low-bush cranberry plant height (mm) (A) and leaf count (B) at 184 days since planting. Above ground (C) and below ground (D) biomass after 1.5 years since planting. Boxplots represent median, and the upper and lower quartiles. Yellow circles represent mean value for each group. Tukey HSD post-hoc comparisons are denoted with lowercase letters above boxplots.

#### Labrador Tea

Labrador tea plants displayed nearly identical trends to both bog blueberry and low-bush cranberry when grown in soils inoculated with microbial communities from the most disturbed treatment site (Figure 4). At the time of final measurements, the mean height and leaf count of Labrador tea grown in the MD treatment was significantly smaller compared to low-bush cranberry grown SD, UD, or ST treatments (Table S5). There was no observed difference between Labrador tea grown in UD, SD, or ST treatments. Labrador tea above ground biomass showed that MD plants weighed in at a significantly smaller amount than plants grown in soils inoculated with UD or SD treatments. No differences were observed between below ground biomass measures (Table S8).

**Figure 4.**
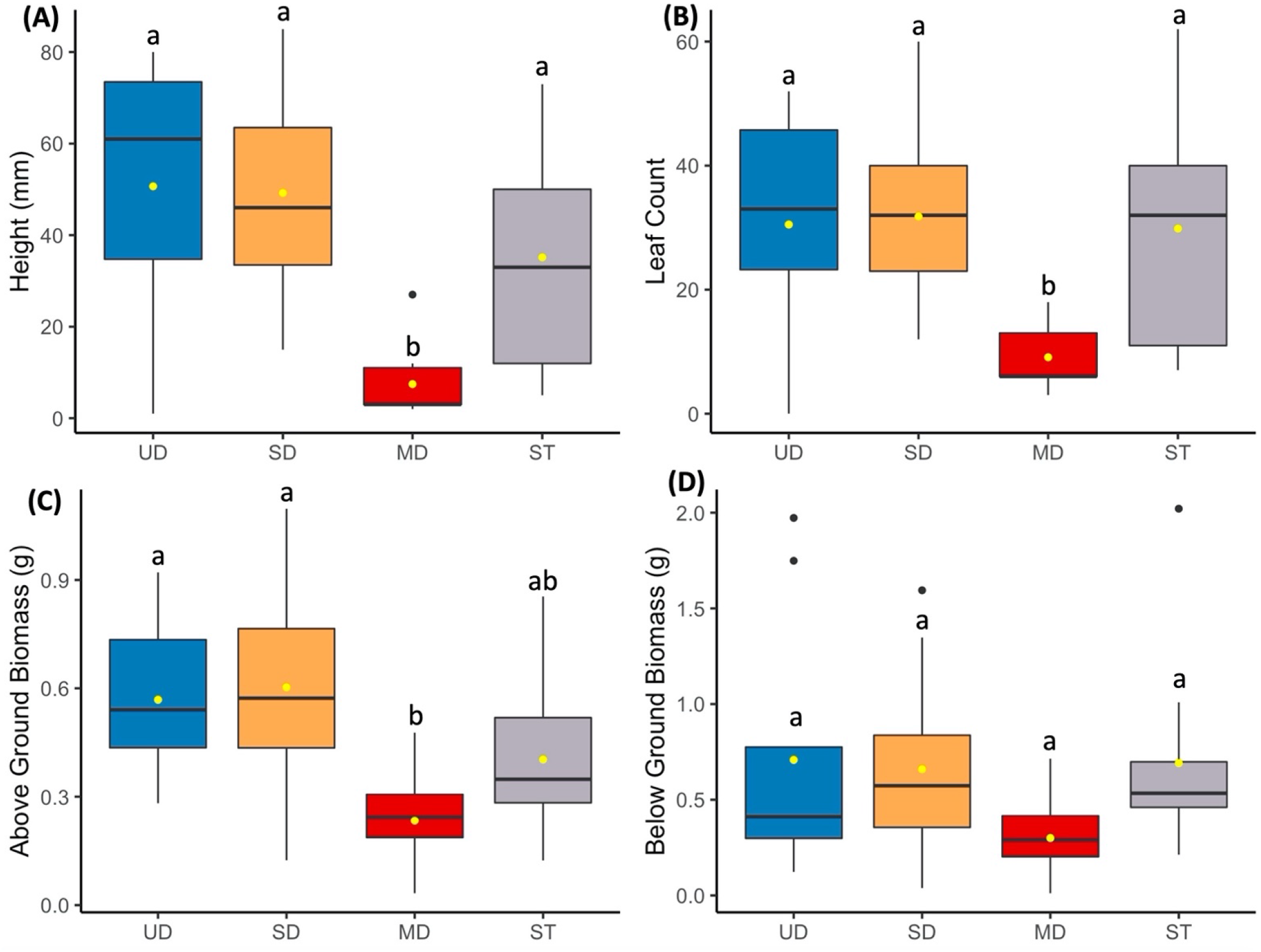
Labrador tea plant height (mm) (A) and leaf count (B) at 184 days since planting. Above ground (C) and below ground (D) biomass after 1.5 years since planting. Boxplots represent median, and the upper and lower quartiles. Yellow circles represent mean value for each group. Tukey HSD post-hoc comparisons are denoted with lowercase letters above boxplots.

#### Fireweed

In contrast to cranberry, bog blueberry, and Labrador tea, fireweed did not show a significant difference in mean leaf count or height (Figure 5); however, fireweed plants grown in inoculant from MD site showed a significant decrease in above ground biomass compared to those grown in UD, SD, or ST treatment soils (Figure 5; Table S6, Table S8).

**Figure 5.**
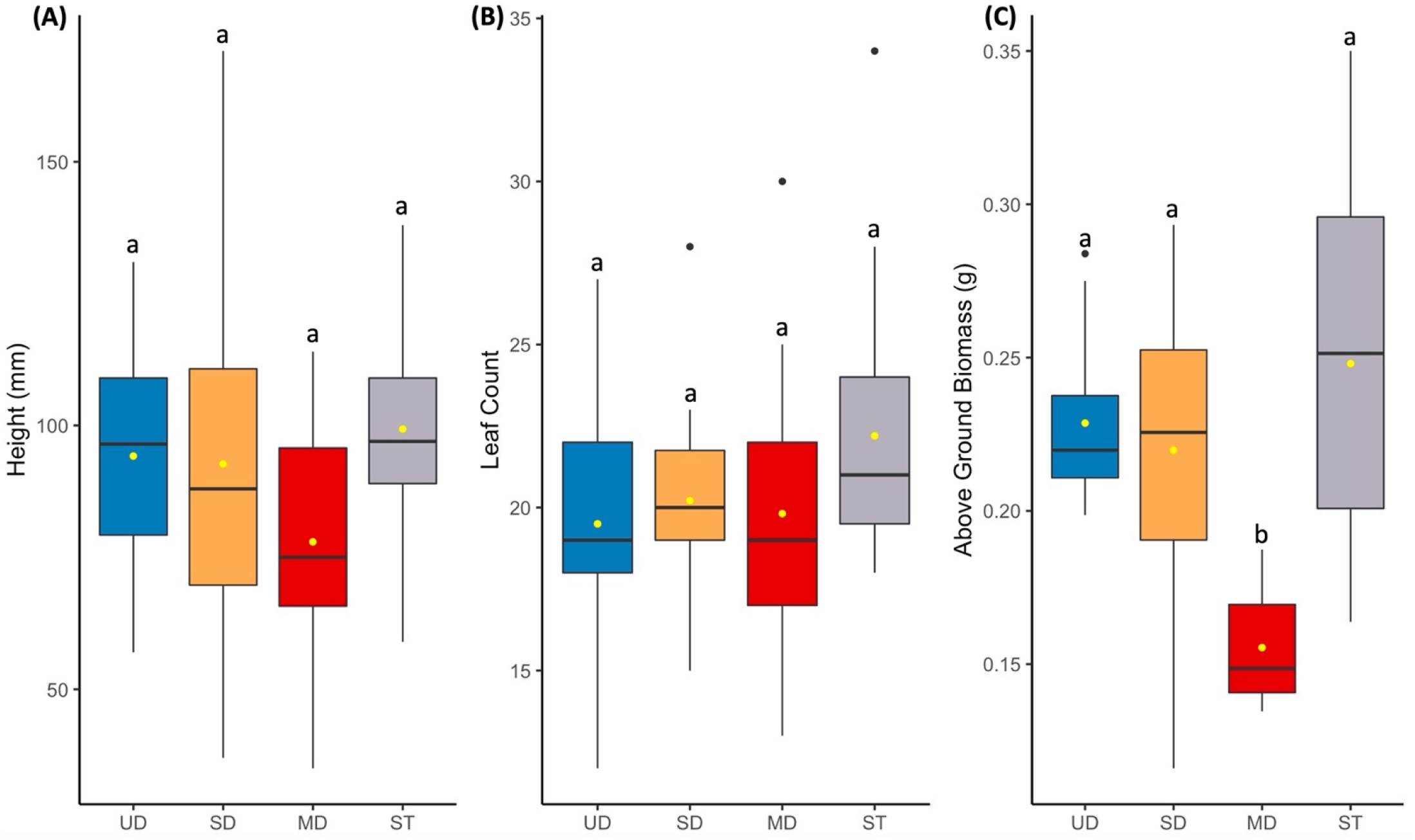
Fireweed plant height (mm) (A) and leaf count (B) at 121 days since planting. Above ground (C) biomass of plants after 165 days since planting. Boxplots represent median, and the upper and lower quartiles. Yellow circles represent mean value for each group. Tukey HSD post-hoc comparisons are denoted with lowercase letters above boxplots.

#### Black Spruce

Black spruce showed no consistent response patterns to microbial inoculant for growth and biomass measures (Figure 6; Table S7). Black spruce plants grew significantly taller when grown with the sterile inoculant (ST) compared to UD, SD, or MD inoculants. Below ground and above ground biomass showed both showed a similar pattern of ST plants measuring at a significantly lower biomass compared to SD plants, and with MD and UD plants showing no significant differences in mass. For leaf (needle) count, no significant differences were observed across all treatment inoculant groups. Leaf count was discontinued and excluded from analysis after needle numbers exceeded 200 for all plants and was not able to be accurately measured.

**Figure 6.**
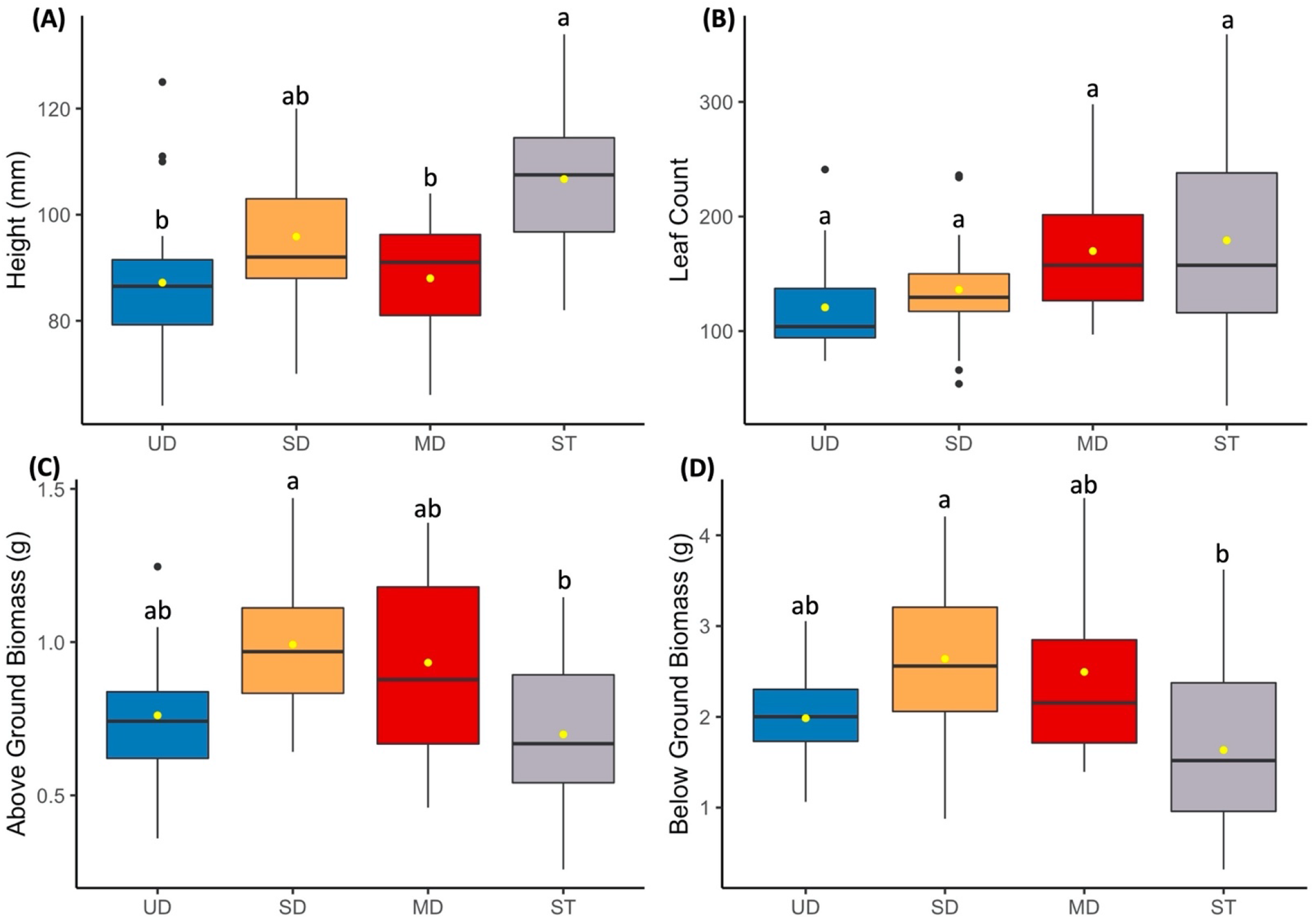
Black spruce height (mm) (A) and leaf count (B) at 184 and 121 days since planting, respectively. Above ground (C) and below ground (D) biomass after 1.5 years since planting. Boxplots represent median, and the upper and lower quartiles. Yellow circles represent mean value for each group. Tukey HSD post-hoc comparisons are denoted with lowercase letters above boxplots.

### 3.2 Microbial Community Analysis

We sequenced 48 metagenomes through four multiplexed MinION runs, and after demultiplexing, total sample reads showed a mean read length of 2594bp, a N50 of 5,531bp, and a mean quality score of 13 (Table S2).

The analysis using Kraken 2 and then Bracken resulted in a mean of 41% (range, 30-60%) of reads being classified as bacterial (60,034 reads). The percent classified did not show a correlation to the read depth of the sample (Figure S3). Of those classified reads, we identified the 10 most common bacterial families across all samples, based on normalized relative abundance, to be: *Acidobacteriaceae, Alcaligenaceae, Bradyrhizobiaceae, Burkholderiaceae, Comamonadaceae, Enterobacteriaceae, Mycobacteriaceae, Pseudomonadaceae, Sphingomonadaceae, Streptomycetaceae* (Figure 7A).

**Figure 7.**
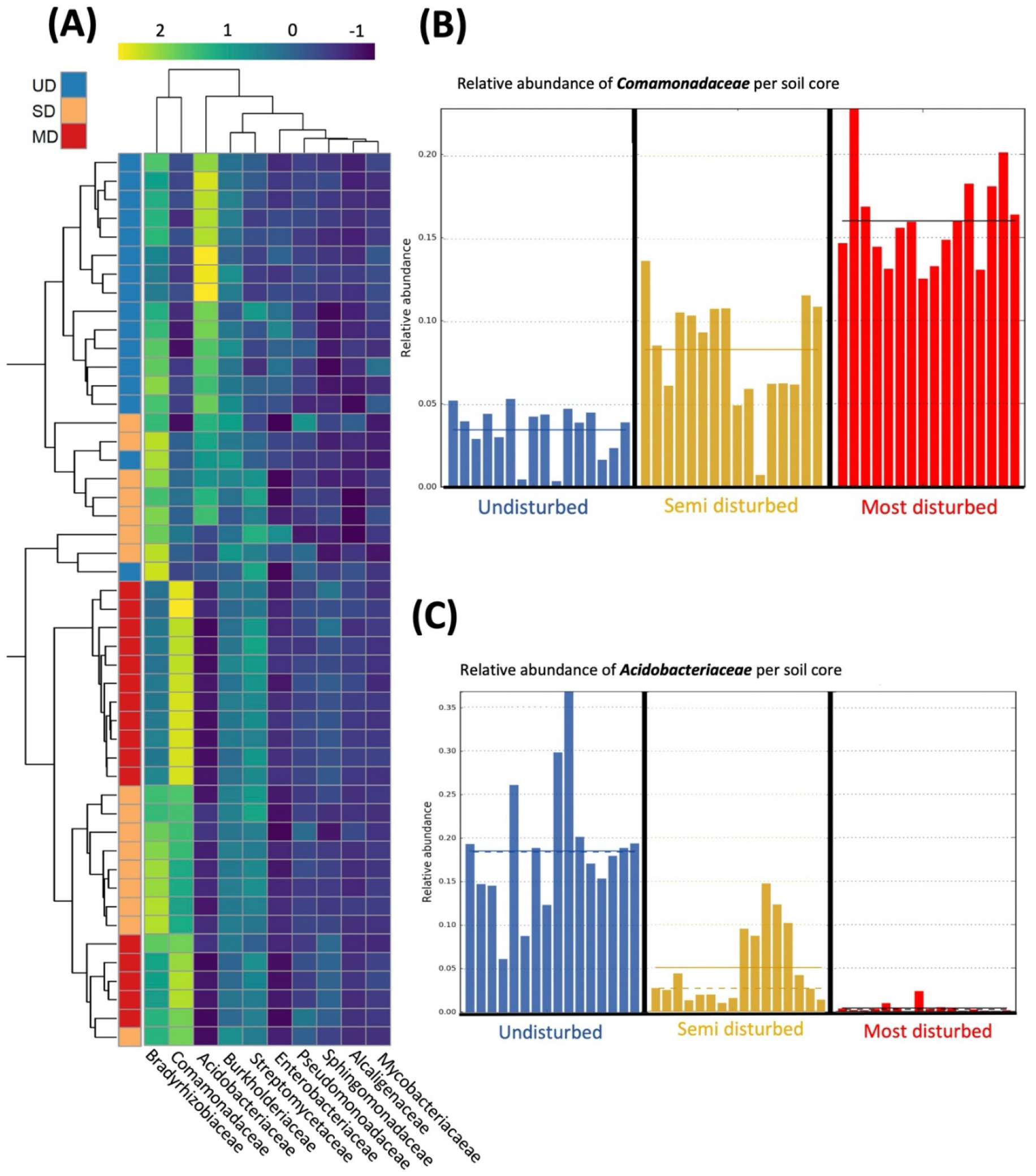
(A) Top 10 most abundant bacterial families classified by Kraken2, scaled by z-score. Samples are clustered by Euclidean distance. (B) Relative abundance of bacterial family, *Comamonadaceae,* per soil core. Solid line represents mean relative abundance per treatment group. (C) Relative abundance of bacterial family, *Acidobacteriaceae,* per soil core. Solid line represents mean relative abundance per treatment group.

Following classification via Kraken 2 and Bracken we utilized the LEfSe method to identify any biomarkers across samples. We observed a general shift in community membership and abundance across the permafrost thaw gradient, with both the UD and MD soil cores showing variation in the dominant families present. Out of only the bacterial reads, the family *Acidobacteriaceae* was found to be significantly enriched in the UD soil microbial communities, present at a mean relative abundance of 18% compared to the SD (5%) and MD (>5%) soil communities (Figure 7C). The family *Comamonadaceae* was found to be present at significantly higher relative abundances within the MD soil microbial communities with a mean of 16%, compared to in UD (5%) and SD (8%) soils (Figure 7B).

## 4 Discussion

Our results indicate that there is a strong relationship between microbial community inoculant and total plant growth. We have evidence that underlying permafrost thaw negatively affects plant growth within several boreal plant species. Within our study we see similar growth response patterns across low-bush cranberry, bog blueberry, Labrador tea, and fireweed plants. Our analyses also revealed that microbial community membership and abundance shifts across active layer soils above a permafrost thaw gradient. Previous research completed by Schütte et al. in 2019 has identified links between microbial communities associated with different degrees of permafrost thaw and plant growth.

Our research showed that plants including bog blueberry, low-bush cranberry, and Labrador tea demonstrated decreased height and leaf count when grown with microbial communities from highly disturbed soil that was associated with deep permafrost thaw compared to undisturbed soil associated with little to no permafrost thaw. Fireweed plants did not show any changes in growth dependent on soil inoculant when looking at leaf count and height. However, when looking at above-ground biomass, plants inoculated with MD soil communities did exhibit lower levels of growth compared to plants inoculated with UD or SD soils. The evidence of this pattern when looking at biomass could potentially be due to the large variation in leaf area and stem thickness that was not present within other plant species that we grew.

We can attribute the changes in plant growth to the variation in microbial communities across the active layer of soil above the permafrost thaw gradient. We hypothesized that if the soil microbial communities were to change depending on the level of associated permafrost thaw, so would the growth of plants in that soil. Consistent with our hypothesis, plants (specifically bog blueberry, low-bush cranberry, Labrador tea, and fireweed) grown with microbial inoculum from MD soils exhibited lower growth than plants grown in soils with no added microbes or with inoculants from the UD or SD soils. These results were similar to a study performed on boreal plant species using microbial inoculums from thermokarst bogs (disturbed soils) and permafrost plateaus (undisturbed soil). Schütte et al. (2019) showed that bog blueberry and marsh-cinquefoil (*Potentilla palustris*) grew significantly worse when inoculated with soil microbes from thermokarst bogs than when inoculated by permafrost plateau soil microbes.

We found no apparent relationship between black spruce growth and the initial soil microbial inoculant. These results correspond to a study (Sniderhan and Baltzer, 2016) analyzing the growth of black spruce across a lateral permafrost thaw gradient in Scotty Creek, Canada, which found that the lateral thaw rate of permafrost did not appear to be a driver of black spruce growth dynamics. Controlled warming experiments meant to simulate the quickly warming northern latitudes have shown that black spruce shoot length tends to increase with warming air and soil temperatures (Bronson et al., 2009; Bronson and Gower, 2010). This suggests that abiotic factors may be having larger influences on black spruce growth compared to biotic factors during climate warming events.

Plant-microbe interactions within the rhizosphere are known determinants of plant health and productivity, and consequently, plant growth (Bever et al., 2010; Reynolds et al., 2003). Interactions between microbes and plants are critical for the acquisition and cycling of nutrients such as biological nitrogen fixation and phosphate solubilization (de Souza et al., 2015). Not only do beneficial microbes assist in nutrient accessibility, but they can also improve plant performance in a variety of systems by conferring resistance to pathogens. Highly studied environments such as agricultural production have shown evidence that small changes to soil microbiomes largely determine the success of plant growth and development (Van Wees et al., 2008; Lutenberg and Kamilova, 2009). In 2019, Wei et al. showed that within an agricultural monoculture system, the composition of initial soil microbial inoculant predetermined whether a crop succumbed to disease or survived. By the end of their growth experiment, microbial communities of healthy plants showed much higher abundances of Acidobacteria than the microbiomes of diseased plants. This is consistent with our finding of the over-representation of *Acidobacteriaceae* in the microbiomes of UD soil.

Variation in plant growth across treatments could largely be due to a change in representation from growth-promoting bacterial and fungal species to deleterious, pathogenic ones found in the soil microbiomes between treatment groups. Beneficial microbes in soil are known to enhance nutrient availability to plants, allowing for an increase in plant growth and productivity (Vimal et al., 2017). Common indicators of a neutral or healthy soil microbiome include wide array of bacterial phyla such as *Proteobacteria, Acidobacteria, Actinobacteria,* and *Bacteriodetes,* as well as and fungal phyla such as *Glomeromycota* (Fierer et al., 2005; Ohsowski et al,. 2014). Consistent with such a roll and perhaps the most striking difference in microbial taxa between treatment groups was the bacterial family *Acidobactericae,* which had a relative abundance of >7% in all UD cores, compared to <0.5% in all MD cores. Members of the *Acidobacteriacae* are thought to be plant-promoters and degrade a variety of simple carbon compounds, such as those that are found in root exudates, which contributes to nutrient cycling and the support of a healthy rhizosphere (Kielak et al., 2016; Campbell, et al., 2010). Other groups of bacteria that were found to be over-represented in the UD soil cores as well, including *Bacillales* which can form endospores and provide protection against and *Enterobacterales*, are both ubiquitous in soil microbiomes.

In contrast, the MD soil cores displayed a much higher relative abundance of bacteria from the order *Comamonadaceae,* a subclass of *Proteobacteria,* than either the UD or SD cores. Members of the order *Comamonadaceae* are known to exhibit pathogenic effects on a variety of plants, including in agricultural settings (Willems et al., 1991). It is possible that the higher presence of groups including known plant-pathogenic bacteria found in the MD soil microbiomes is leading to the decrease in associated plant growth, caused by a disruption in nutrient cycling and direct alterations to the plant rhizospheres. It is important to note that our results suggest that a decrease in plant growth can be linked to changes in the taxonomic diversity of microbial communities, but further experimentation investigating the functional genetic potential of the microbial communities will be required to elucidate the mechanisms that underlie the patterns described here.

Having an increased understanding of how microbial communities that reside above permafrost affect plant growth is important for predicting effects of permafrost thaw on plant communities and ecosystem health. Specifically, for potentially predicting effects on plants, such as bog blueberry, low-bush cranberry, and Labrador tea, that are relied on as common food sources for both humans and animals throughout the Northern regions. Although the links between soil microbial and plant communities are complex and rarely straight-forward, this study indicates that the variation in active layer soil microbiomes can have a strong effect on plant growth. This data point towards building an understanding of how shifting microbial communities may affect plant growth in the face of climate change causing thawing permafrost; specifically, growth of plants such as bog blueberry and low-bush cranberry, both plants integral to the diets of many Alaska Native cultures. The results reported here demonstrate that soil microbes have the capability to alter plant growth in response to climate change, therefore highlighting the need for further research on the taxonomy and functionality of microbial communities residing above thawing permafrost.

## Supporting information

SFigure 1

## 5 Acknowledgements

We would like to thank Dr. Anne-Lise Ducluzeau and Tracie Haan for laboratory support, and Scout McDougall, Jennie Humphrey for support with data collection. We would also like to thank the invaluable support in the greenhouse from Drs. Mel Durret and Mark Wright. Thank you to Dr. Tom Douglas from the Cold Regions Research and Engineering Laboratory (CRREL)-Alaska for providing access to the field site. This research was supported by the Alaska BLaST program, the Institute of Arctic Biology, and Alaska INBRE.

## 6 Author Contributions

US and DD contributed to the conception and design of the study. TS performed the data collection and statistical analysis and wrote the first draft of the manuscript. All authors contributed to the manuscript revisions and approved the submitted version.

## 8 Data Availability Statement

The sequencing data discussed in this manuscript have been deposited on EBI Metagenomics, accession numbers xxx.

