## Supplementary material for "Soil disturbance affects plant growth via soil microbial community shifts": SFigure 1

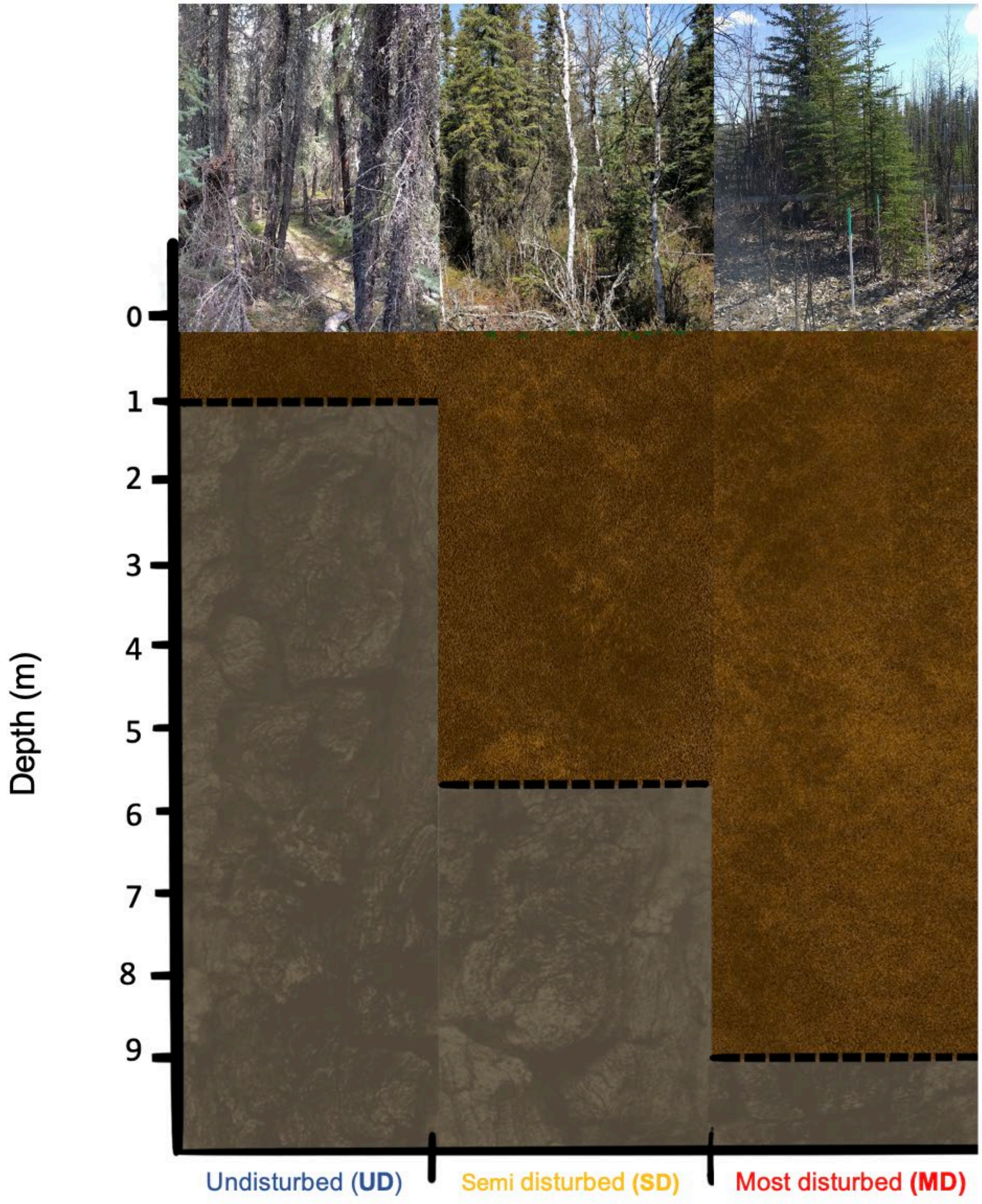


**Figure S1.** Fairbanks Permafrost Experiment Station (FPES) plot layout and permafrost thaw levels and above ground plant cover

| Run | Flow cell + kit model |
|---|---|
| 20180611 | dna_r9.5_450bps |
| 20180719A | dna_r9.4.1_450bps_hac |
| 20180719B | dna_r9.4.1_450bps_hac |
| 20180719C | dna_r9.4.1_450bps_hac |

**Table S1.** Base calling specifications per sequencing run

**Table S2.** Summary of sequencing statistics per sample following quality control

| Sample | Run | Barcode | Total yield (bp) | Total read count (bp) | Avg read length (bp) |
|---|---|---|---|---|---|
| MD 9.1 | FPES_20180719A | 05 | 400,202,630 | 194,937 | 2052.98 |
| MD 9.2 | FPES_20180719B | 06 | 309,392,310 | 107,312 | 2883.11 |
| MD 9.3 | FPES_20180611 | 09 | 203,489,778 | 93,380 | 2179.16 |
| MD 9.4 | FPES_20180719C | 05 | 373,793,563 | 140,143 | 2667.23 |
| MD 10.1 | FPES_20180719A | 08 | 335,540,391 | 107,848 | 3111.23 |
| MD 10.2 | FPES_20180611 | 10 | 237,477,806 | 66,156 | 3589.66 |
| MD 10.3 | FPES_20180719B | 07 | 266,280,833 | 95,477 | 2788.95 |
| MD 10.4 | FPES_20180719C | 06 | 528,466,953 | 199,447 | 2649.66 |
| MD 11.1 | FPES_20180611 | 11 | 339,699,261 | 141,202 | 2405.77 |
| MD 11.2 | FPES_20180719A | 07 | 319,858,083 | 151,923 | 2105.4 |
| MD 11.3 | FPES_20180719B | 05 | 244,526,616 | 87,798 | 2785.1 |
| MD 11.4 | FPES_20180719C | 03 | 464,942,414 | 152,761 | 3043.59 |
| MD 12.1 | FPES_20180719A | 01 | 242,429,913 | 104,932 | 2310.35 |
| MD 12.2 | FPES_20180611 | 12 | 187,202,338 | 87,766 | 2132.97 |
| MD 12.3 | FPES_20180719B | 04 | 237,416,454 | 134,994 | 1758.72 |
| MD 12.4 | FPES_20180719C | 02 | 626,845,953 | 198,765 | 3153.7 |
| SD 9.1 | FPES_20180719A | 04 | 683,763,421 | 392,010 | 1744.25 |
| SD 9.2 | FPES_20180719B | 01 | 400,235,372 | 173,176 | 2311.15 |
| SD 9.3 | FPES_20180719C | 07 | 737,067,162 | 299,103 | 2464.26 |
| SD 9.4 | FPES_20180611 | 05 | 203,634,783 | 77,269 | 2635.4 |
| SD 10.1 | FPES_20180719A | 03 | 435,414,246 | 203,659 | 2137.96 |
| SD 10.2 | FPES_20180719B | 03 | 308,506,891 | 136,862 | 2254.15 |
| SD 10.3 | FPES_20180611 | 06 | 304,500,863 | 101,986 | 2985.71 |
| SD 10.4 | FPES_20180719C | 12 | 465,408,883 | 140,711 | 3307.55 |
| SD 11.1 | FPES_20180719A | 02 | 269,690,205 | 129,469 | 2083.05 |
| SD 11.2 | FPES_20180611 | 07 | 182,991,407 | 53,543 | 3417.65 |
| SD 11.3 | FPES_20180719B | 12 | 264,072,945 | 100,301 | 2632.8 |
| SD 11.4 | FPES_20180719C | 10 | 651,807,046 | 220,446 | 2956.77 |
| SD 12.1 | FPES_20180719A | 06 | 523,124,325 | 263,679 | 1983.94 |
| SD 12.2 | FPES_20180611 | 08 | 345,914,971 | 169,926 | 2035.68 |
| SD 12.3 | FPES_20180719B | 02 | 486,981,032 | 246,027 | 1979.38 |
| SD 12.4 | FPES_20180719C | 08 | 699,809,340 | 184,567 | 3791.63 |
| UD 9.1 | FPES_20180611 | 01 | 235,098,257 | 80,943 | 2904.49 |
| UD 9.2 | FPES_20180719A | 09 | 335,750,695 | 122,957 | 2730.64 |
| UD 9.3 | FPES_20180719B | 09 | 316,380,483 | 134,144 | 2358.51 |
| UD 9.4 | FPES_20180719C | 01 | 450,701,122 | 147,992 | 3045.44 |
| UD 10.1 | FPES_20180719A | 10 | 385,328,155 | 124,052 | 3106.18 |
| UD 10.2 | FPES_20180611 | 02 | 275,828,426 | 78,460 | 3515.53 |
| UD 10.3 | FPES_20180719B | 08 | 296,873,474 | 106,709 | 2782.08 |
| UD 10.4 | FPES_20180719C | 04 | 556,101,883 | 178,135 | 3121.8 |

| | | | | | |
|---|---|---|---|---|---|
| UD 11.1 | FPES_20180611 | 03 | 295,277,023 | 97,759 | 3020.46 |
| UD 11.2 | FPES_20180719A | 11 | 344,805,302 | 116,453 | 2960.9 |
| UD 11.3 | FPES_20180719B | 11 | 206,576,878 | 101,380 | 2037.65 |
| UD 11.4 | FPES_20180719C | 11 | 516,115,731 | 135,427 | 3811.03 |
| UD 12.1 | FPES_20180719A | 12 | 272,044,502 | 108,655 | 2503.75 |
| UD 12.2 | FPES_20180611 | 04 | 316,589,380 | 90,290 | 3506.36 |
| UD 12.3 | FPES_20180719B | 10 | 293,661,051 | 121,689 | 2413.21 |
| UD 12.4 | FPES_20180719C | 09 | 546,467,017 | 207,151 | 2638.01 |

**Table S3.** ANOVA results for bog blueberry growth measures

| Growth Measure | Source | Degrees of freedom | Sum of squares | Mean sum of squares | F value | P value |
|---|---|---|---|---|---|---|
| Height | FPES | 3 | 52965 | 17655 | 11.44 | **$5.88 \times 10^{-6}$** |
| | Residuals | 56 | 86395 | 1543 | | |
| Leaf Count | FPES | 3 | 16568 | 5523 | 5.544 | **0.00207** |
| | Residuals | 57 | 56778 | 996 | | |
| Above Ground Biomass | FPES | 3 | 0.8878 | 0.2959 | 6.663 | **0.000787** |
| | Residuals | 46 | 2.043 | 0.04441 | | |
| Below Ground Biomass | FPES | 3 | 0.926 | 0.30862 | 4.189 | **0.0106** |
| | Residuals | 46 | 3.389 | 0.07367 | | |

*Bolded p-value indicates significance with a < 0.05

**Table S4.** ANOVA results for low-bush cranberry growth measures

| Growth Measure | Response | Degrees of freedom | Sum of squares | Mean sum of squares | F value | P value |
|---|---|---|---|---|---|---|
| Height | FPES | 3 | 13881 | 4627 | 8.966 | **$6.25 \times 10^{-5}$** |
| | Residuals | 55 | 28385 | 516 | | |
| Leaf Count | FPES | 3 | 6502 | 2167.3 | 9.295 | **$4.39 \times 10^{-5}$** |
| | Residuals | 56 | 13057 | 233.2 | | |
| Above Ground Biomass | FPES | 3 | 1.852 | 0.6173 | 12.98 | **$2.32 \times 10^{-6}$** |
| | Residuals | 49 | 2.331 | 0.0476 | | |
| Below Ground Biomass | FPES | 3 | 0.525 | 0.17516 | 2.019 | 0.124 |
| | Residuals | 49 | 4.252 | 0.08677 | | |

*Bolded p-value indicates significance with a < 0.05

**Table S5.** ANOVA results for Labrador tea growth measures

| Growth Measure | Response | Degrees of freedom | Sum of squares | Mean sum of squares | F value | P value |
|---|---|---|---|---|---|---|
| Height | FPES | 3 | 12314 | 4105 | 7.947 | 0.000233 |
| | Residuals | 45 | 23243 | 517 | | |
| Leaf Count | FPES | 3 | 3476 | 1158.7 | 5.216 | 0.00355 |
| | Residuals | 45 | 9995 | 222.1 | | |
| Above Ground Biomass | FPES | 3 | 0.8639 | 0.28795 | 5.65 | 0.00273 |
| | Residuals | 37 | 1.8858 | 0.0509 | | |
| Below Ground Biomass | FPES | 3 | 6.6 | 2.1994 | 2.377 | 0.0855 |
| | Residuals | 37 | 34.23 | 0.9251 | | |

*Bolded p-value indicates significance with a < 0.05

**Table S6.** ANOVA results for fireweed growth measures

| Growth Measure | Response | Degrees of freedom | Sum of squares | Mean sum of squares | F value | P value |
|---|---|---|---|---|---|---|
| Height | FPES | 3 | 13439 | 4480 | 3.053 | **0.0356** |
| | Residuals | 57 | 83628 | 1467 | | |
| Leaf Count | FPES | 3 | 0.1567 | 0.05224 | 1.447 | 0.239 |
| | Residuals | 55 | 1.9858 | 0.0361 | | |
| Above Ground Biomass | FPES | 3 | 1.51 | 0.5034 | 11.17 | **$7.24 \times 10^{-6}$** |
| | Residuals | 57 | 2.57 | 0.0451 | | |

*Bolded p-value indicates significance with a < 0.05

**Table S7.** ANOVA results for black spruce growth measures

| Growth Measure | Response | Degrees of freedom | Sum of squares | Mean sum of squares | F value | P value |
|---|---|---|---|---|---|---|
| **Height** | FPES | 3 | 3634 | 1211.3 | 5.654 | **0.00186** |
| | Residuals | 57 | 12231 | 214.6 | | |
| **Leaf Count** | FPES | 3 | 1.292 | 0.4307 | 2.378 | 0.0787 |
| | Residuals | 60 | 10.869 | 0.1811 | | |
| **Above Ground Biomass** | FPES | 3 | 0.871 | 0.29023 | 4.154 | **0.0101** |
| | Residuals | 55 | 3.843 | 0.06987 | | |
| **Below Ground Biomass** | FPES | 3 | 9.71 | 3.236 | 4.264 | **0.00896** |
| | Residuals | 54 | 40.99 | 0.759 | | |

*Bolded p-value indicates significance with a < 0.05

Height for all plants was compared at day 184, after the growth rate had started to slow (except fireweed which was compared at day 121 due to harvest for biomass). Leaf count was compared at day 184 for BB, CB, and LT after growth had stabilized. Leaf count for fireweed was analyzed on day 121 due to harvest, and black spruce was analyzed on day 121 (the last needle count collected, due to challenges with collection accuracy). Only above ground biomass was collected for fireweed on day 121. Above ground biomass was collected 640 days following planting for all other plant types (BB, CB, BS, and LT). Below ground biomass was collected at a delayed date 665 days after planting, due to complications related to COVID-19.

**Table S8.** Tukey's honestly significant difference (HSD) post-hoc test results per growth measure and plant type

| | GROWTH MEASURE | UD | SD | MD | ST |
|---|---|---|---|---|---|
| **Blueberry** | Height | a | a | b | a |
| | Leaf Count | a | a | b | a |
| | Above Ground Biomass | ab | a | b | b |
| | Below Ground Biomass | ab | a | ab | b |
| **Cranberry** | Height | a | a | b | a |
| | Leaf Count | a | a | b | a |
| | Above Ground Biomass | b | a | b | b |
| | Below Ground Biomass | a | a | a | a |
| **Labrador tea** | Height | a | a | b | a |
| | Leaf Count | a | a | b | a |
| | Above Ground Biomass | a | a | b | ab |
| | Below Ground Biomass | a | a | a | a |
| **Fireweed** | Height | ab | a | ab | b |
| | Leaf Count | a | a | a | a |
| | Above Ground Biomass | a | a | b | a |
| **Black spruce** | Height | b | ab | b | a |
| | Leaf Count | a | a | a | a |
| | Above Ground Biomass | ab | a | ab | b |
| | Below Ground Biomass | ab | a | ab | b |

**Figure S2.** Average height over time for each plant type

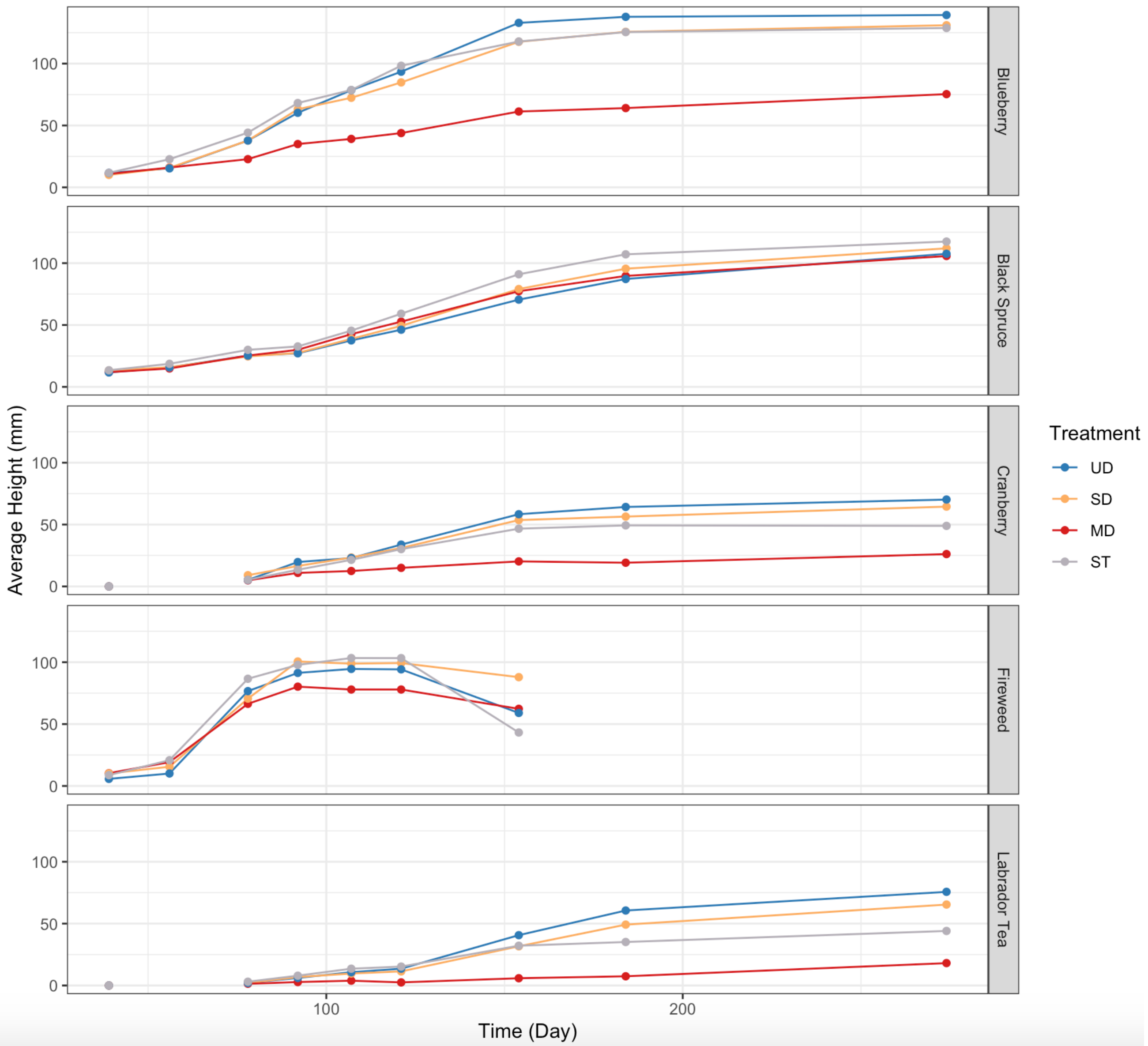


*Fireweed growth ends on day 121 when the plants were harvested for above ground biomass

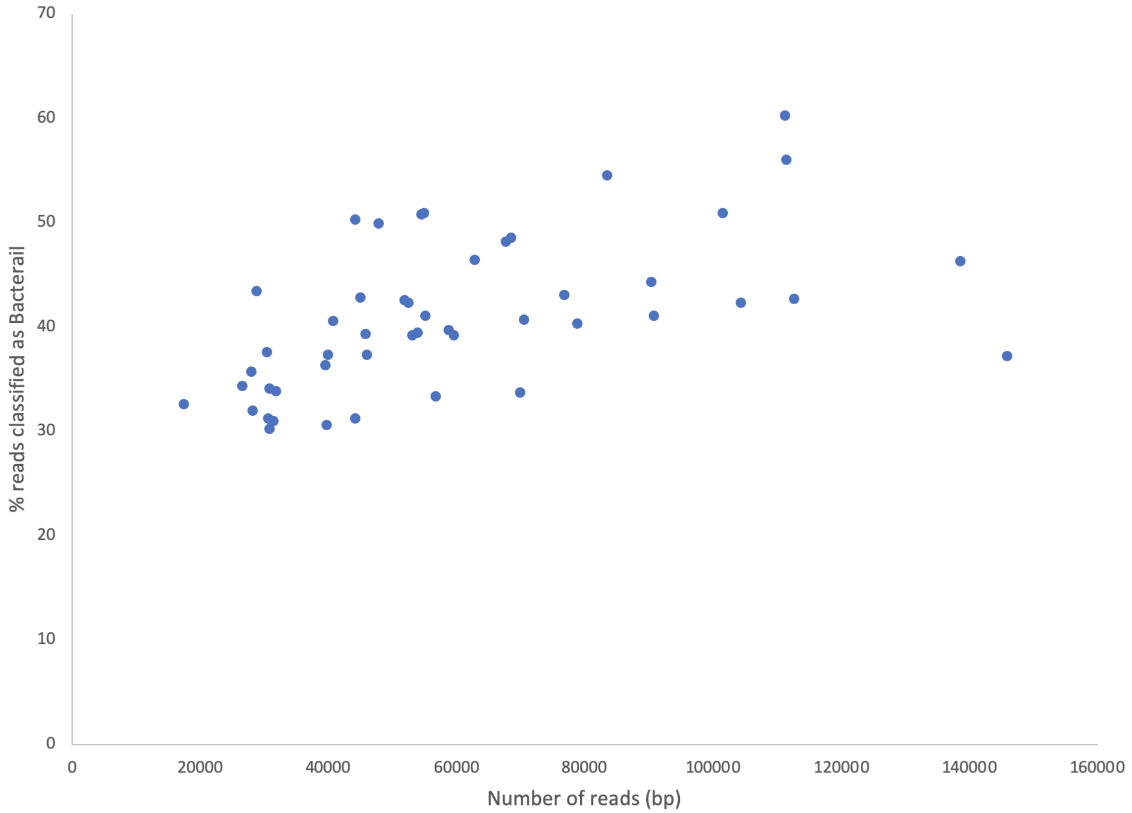


**Figure S3.** Percentage of metagenomic reads classified as bacterial and the number of reads classified via Kraken2 (n = 48)
